# Experience resetting in reinforcement learning facilitates exploration–exploitation transitions during a behavioral task for primates

**DOI:** 10.1101/2021.09.30.462676

**Authors:** Kazuhiro Sakamoto, Hidetake Okuzaki, Akinori Sato, Hajime Mushiake

## Abstract

The exploration–exploitation trade-off is a fundamental problem in reinforcement learning. To study the neural mechanisms involved in this problem, a target search task in which exploration and exploitation phases appear alternately is useful. Monkeys well trained in this task clearly understand that they have entered the exploratory phase and quickly acquire new experiences by resetting their previous experiences. In this study, we used a simple model to show that experience resetting in the exploratory phase improves performance rather than decreasing the greediness of action selection, and we then present a neural network-type model enabling experience resetting.

## 1 Introduction

Accumulating experience and knowledge and then applying them to action decisions is an extremely important foundation for success. However, in order to adapt to a constantly changing environment, it is also essential to reset the past experience and knowledge and search for new strategies. This question of whether to use or discard experience and knowledge, termed the exploration–exploitation trade-off, is one of the fundamental problems of reinforcement learning theory [1].

The target search task is useful for studying the neural mechanisms involved in the exploration–exploitation trade-off [2-4]. In the target search task, when a subject selects the correct target for a certain number of trials in a row, the correct target is switched without instruction, and the subject must search for a new correct target (Fig. 1). In other words, in this task, the exploitation phase alternates with the exploration phase, and the subject’s performance and related neuronal activities can provide clues as to how the brain solves the exploration–exploitation problem. In our previous studies, we showed that well-trained monkeys, after a series of rewarded trials, quickly reset their previous behavior when they get an incorrect outcome due to a switch in the correct target without instruction and immediately start exploratory behavior, finding a new correct target in a small number of trials [2,3]. Furthermore, we found neurons in the supplementary eye field of the medial frontal lobe that encoded the beginning of the exploratory phase. The existence of these cells indicates that animals make a clear distinction between exploratory and exploitatory behaviors [3].

**Fig. 1.**
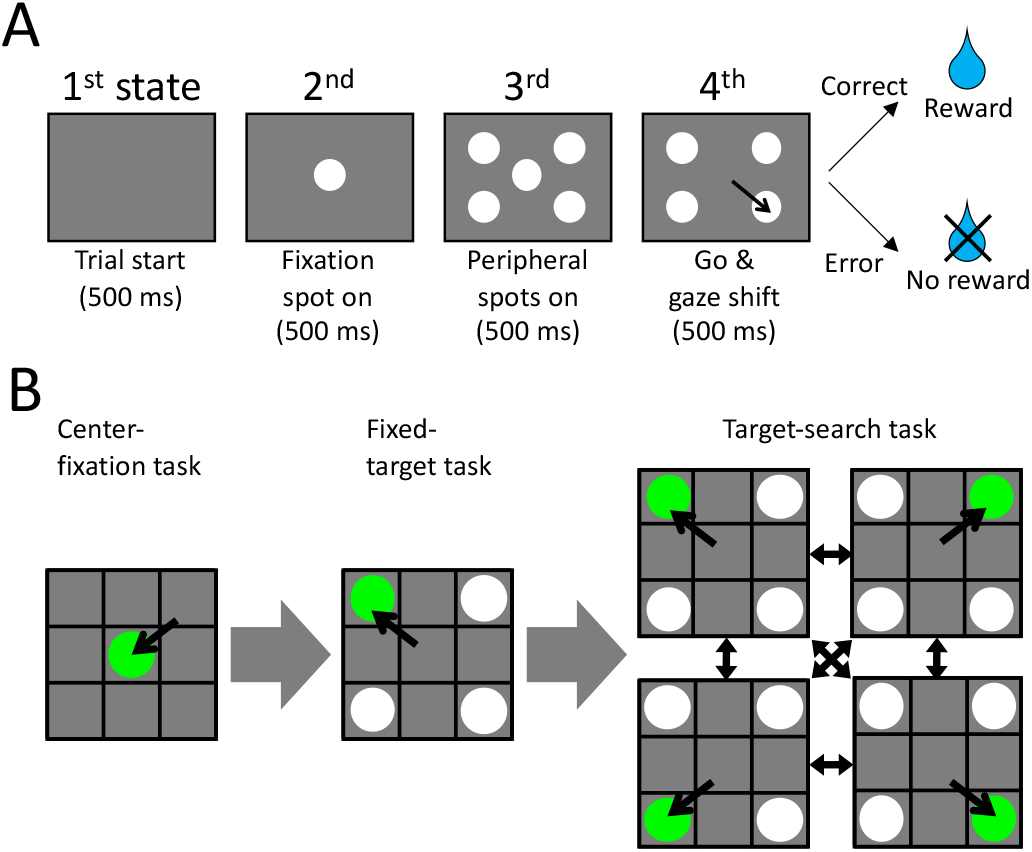
The target search task and pre-lessons. (A) The event sequence of the target search task. (B) Schematics of the pre-lessons of the center-fixation task (left) and fixed-target (middle) tasks. Targets are designated by green color for distinction.

Khamassi et al. (2015) created a reinforcement learning model to analyze neuronal activity in the frontal lobe involved in exploratory and exploitatory behavior [4]. The model quickly shifted to the exploratory behavior mode after the correct target was switched by reducing the greediness in their action selection, *i*.*e*., by reducing the inverse temperature of the softmax function for action selection during the exploratory phase. However, in their model, one time-step of computation is one trial, *i*.*e*., the model does not learn the event sequence of the task. Therefore, the meta-learning of inverse temperature regulation may also impair appropriate behavior, such as maintaining fixation, in each event during one trial.

Here, we show the usefulness of the experience resetting in the quick transition from the exploitation phase to the exploration phase. First, in simple temporal difference (TD) learning models that learn the sequence of events and the responses to them in the target search task, we demonstrate that resetting the value function is a metalearning strategy that leads to better performance than the inverse temperature regulation in the exploratory phase. Based on these results, we present a neural network reinforcement learning (NNRL) model as a step toward building a model that is closer to the brain and can adapt to complex situations in the future. This model quickly transitions to exploratory behavior and acquires behavior for new targets by resetting previous experience, while taking appropriate actions at each event of the task.

## 2 Methods

### 2.1 Behavioral tasks and simulation framework

The target search task includes the four task periods “trial start,” “fixation spot on,” “peripheral spots on,” and “go & gaze shift” (Fig. 1A). During the “fixation spot on” period, the gaze is directed to the central spot (C). In the subsequent “peripheral spots on” period, the subject is required to keep fixating on spot (C) without being disturbed by the four spots presented around it, left-up (LU), right-up (RU), left-down (LD), and right-down (RD). When (C) disappears at the “go & gaze shift” period, the subject shifts his/her gaze to one of the four surrounding spots, and if it is the correct target, he/she is rewarded. If the subject answers correctly for a certain number of trials in a row, the target is switched without any instruction, and the subject is required to search for the new one. The duration of each period in the experiments with primates was 500 ms [2, 3].

The agent required two pre-lessons before learning the target search task (Fig. 1B). The first one was a center-fixation task. This task included only two periods, “trial start” and “fixation spot on,” and a reward was given for viewing (C) during the “fixation spot on” period. In the second step, the agent started to learn the fixed-target task. This task involves the same four periods as the target search task, but a particular target continues to be the correct target. Importantly, the agent had to be trained on the four targets in an unbiased order at this stage. In the following simulations, the time step for calculation and the number of consecutive correct trials for target switch were fixed at one task period and 15 trials, respectively.

### 2.2 Simple TD learning models with meta-learning

First, we built a conventional TD learning model that makes it easy to understand what changes are occurring during the learning process. Assuming that the four states *S* of the presentation screen, *i*.*e*., the four task periods (Fig. 1A), are known, we explicitly implemented them to be distinguished. By differentiating the screen into three by three compartments, we established that the agent can choose nine actions *A, i*.*e*., gaze to positions left-up (LU), up (U), right-up (RU), left (L), center (C), right (R), left-down (LD), down (D) and right-down (RD) (see also Fig. 2). Therefore, the state value function *V* is given by a 4 × 9 matrix. The initial values were all set to 0.5.

**Fig. 2.**
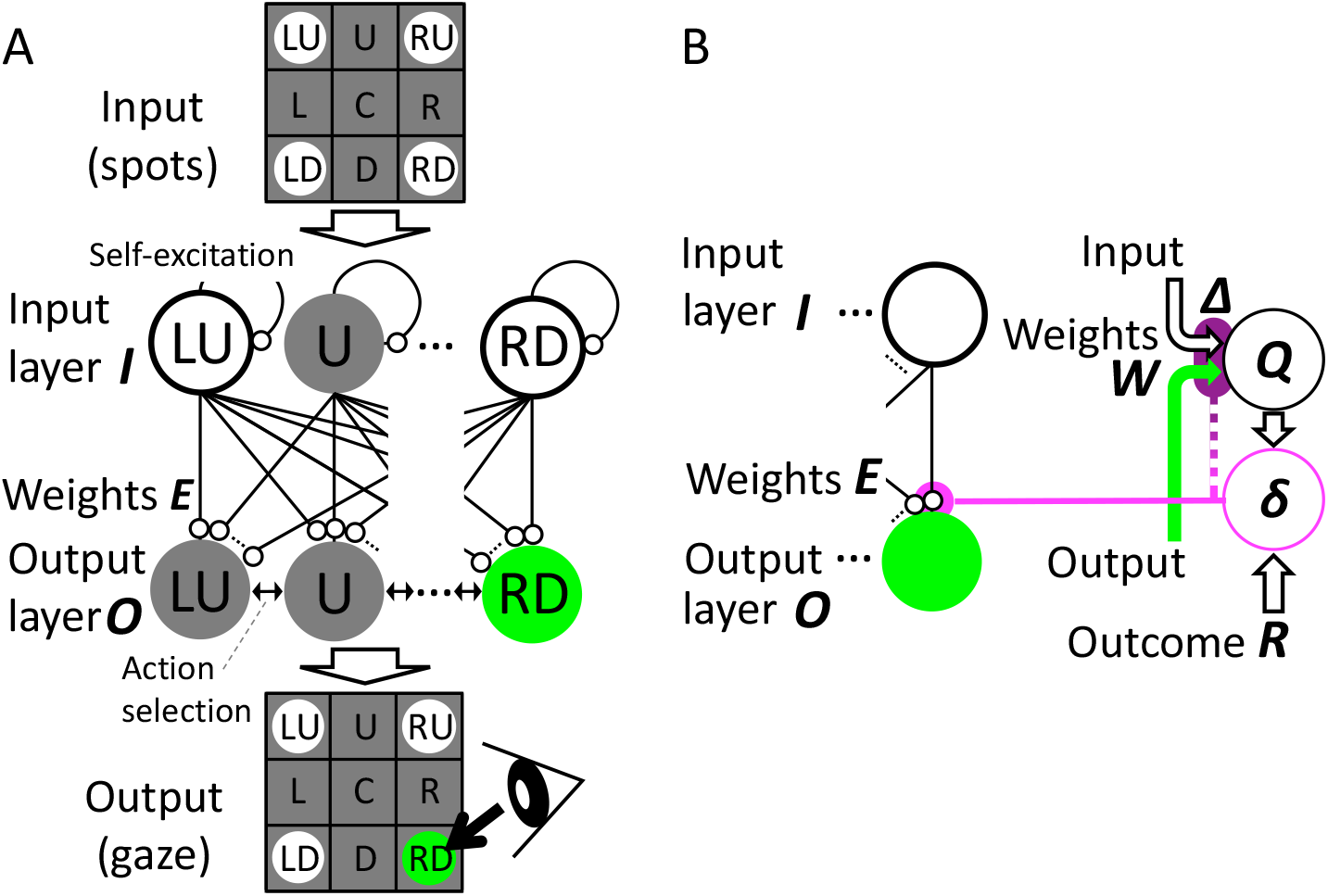
Schematic view of the model. (A) The input–output part. The display was divided into nine areas for simplicity. (B) The reward expectation part. *Δ* and *δ* are different values, although the former is obtained from the latter. Thus, they are connected by a line with color gradation.

The action was selected by the softmax function. That is, the probability that an action *a*_*j*_ is selected at time step *t + 1* is determined by the following equation:

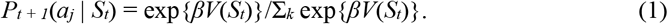

*β* is the inverse temperature. The higher the value of *β* is, the greedier the agent will be; *i*.*e*., the agent will choose only the action with the highest action value, and the lower the value is, the more likely it becomes that each action will be chosen with equal probability. Basically, *β* was set to 100 so that actions were selected in a greedy manner. Using the reward prediction error

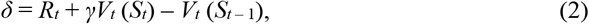

the *V* table is updated according to the following equation:

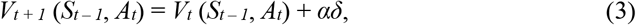

where *R, γ*, and *α* are the reward, the discount rate, and the learning rate, respectively. The default values of *γ* and *α* were set to 0.7 and 0.1, respectively, to get the best results in this standard TD learning model.

We examined the effects of meta-learning, *i*.*e*., regulating parameters other than the *V* value during the exploratory phase for the next two cases. In the first, following Khamassi et al. (2015) [4], we randomized action selection by reducing the inverse temperature. In the second, we reset the reward expectation, *i*.*e*., the corresponding *V*_*t*_(*S*_*t*_), to its initial value. The judgement of the exploratory phase was simply based on the detection of no reward, as the agent’s action selection was so greedy that it made no mistake in the exploitation phase. We also assumed that the task phase in which outcome was given was known, and performed the above two meta-learnings only in the fourth state.

### 2.3 NNRL model

Next, as a model that is closer to the brain and can adapt to complex environments, we constructed a NNRL model. The input–output part of the model (Fig. 2A) has an input layer *I* consisting of nine units corresponding to the vectors *D* of the on–off of the nine spots, and an output layer *O* consisting of nine units corresponding to the actions *A* toward the nine locations. In this model, the four states of spot presentation (Fig. 1A) are assumed to be unknown. However, in order to make it easier to distinguish these four patterns implicitly and to produce a tendency to persistently look at a single point, each unit *I*_*i*_ of the input layer has a self-excitation mechanism, as shown in the second term of the numerator of the following equation:

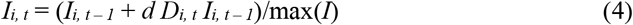

so that the activity of the unit *I*_*i*_ increases relatively when the corresponding spot *D*_*i*_ is continuously presented. The normalization by the maximum value of the input layer serves to keep the activity of the units in the range of 0–1, where *d* is an arbitrary positive decay constant. The activity of each unit in the output layer is determined by the following equation:

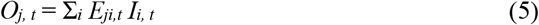

where *E*_*ji*_ is the connection weight from the unit *I*_*i*_ in the input layer to the unit *O*_*j*_ in the output layer.

The mechanism involved in updating the input–output connection weight matrix *E* is schematized in Fig. 2B. First, the action value function matrix *Q*, using both the spot input *D*_*i*_ and the selected action *A*_*j*_ with the coefficient matrix *W*, is calculated as:

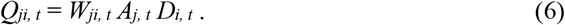

Note that, as a result, the update of the connection weights *E* at each time step is pin-pointed only for the unit *O*_*j*_ corresponding to the selected action *A*_*j*_, as shown in

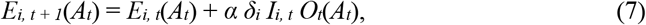

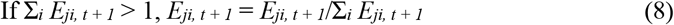

using the prediction error

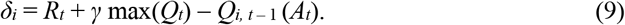

*R* is the reward, *γ* is the discount factor, *α* is the learning rate, and *γ* and *α* were fixed at 0.9 and 0.25, respectively, which yielded good results. The max function was used to extract any predictions of reward, but there are other ways to implement it. As shown in equation (8), if the sum of the input–output connection weights to the unit *O*_*i*_ exceeds 1, each value was normalized by that sum, which contributed to equalizing the connection weights to unit *O*_*i*_. If this equalization is incomplete, *E* will not be completely reset, and as a result, the agent will not switch targets in the target search task.

Meanwhile, the coefficient matrix *W* for the value function *Q* is updated to take only the values −1, 0, and 1 so that the input−output connection weight to *O*_*j*_ corresponding to the selected action *A*_*j*_ is reset at a time, as follows:

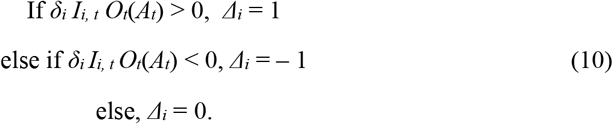

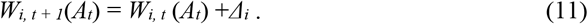

For action selection, a mechanism that allows a winner-take-all outcome by competition with lateral inhibition, *etc*., is desirable as a realistic implementation, but, for simplicity and to compare as closely as possible with the TD learning models above, we used the same softmax function as in equation (1). However, the input is not *Q*, but rather the value of the output layer *I*. When all the values in the output layer were zero, each unit was given a value of 1/9, so that the action would be chosen randomly. Note that the NNRL model does not include explicit mechanisms for search period judgement and meta-learning as in the above TD learning models.

## 3 Results

### 3.1 Investigating the effect of meta-learning on the transition between exploration and exploitation using simple TD learning models

First, to study how learning models with meta-learning for a quick transition from exploitation phase to exploration phase behave, we built TD learning models that learn the target search task (Fig. 1A) and investigated the effects of inverse temperature regulation and experience resetting during the exploration phase. The computational experiments went through the stages shown in Fig. 1B. In all cases, the target search task was not learned before the center-fixation task was performed. After 1,000 trials of the center-fixation task, and 1,000 trials of the fixed-target tasks with each targeting LU, RU, LD, and RD, the models were served to perform 10,000 trials of the target search task, in which the target was switched after 15 consecutive correct trials. The percentage of correct responses and the number of target switches were evaluated; changes in these values with the learning rate are shown in Fig. 3.

**Fig. 3.**
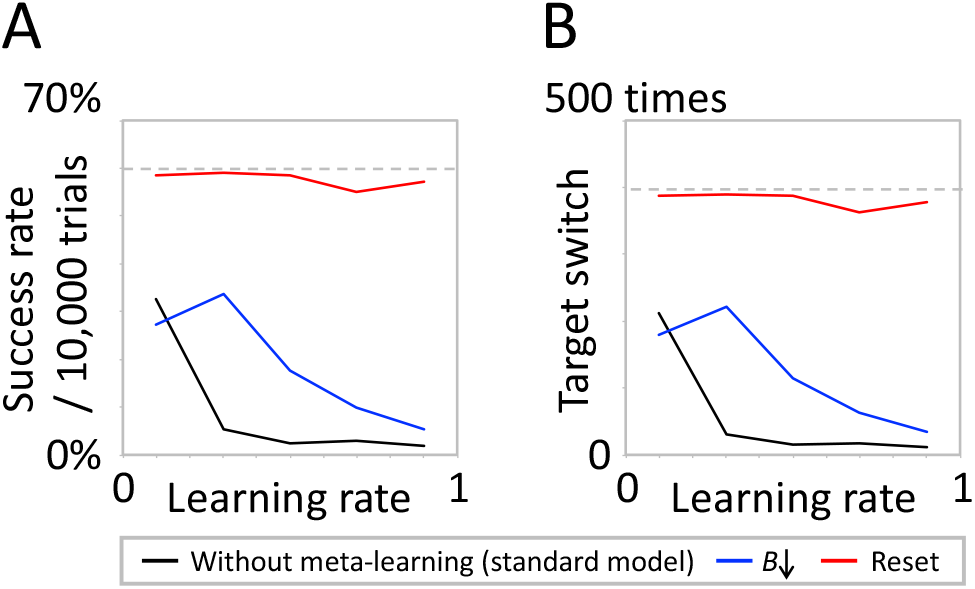
Performances of the TD learning models during a 10,000-trial session after pre-lessons. (A) Success rate. (B) Number of target switches. The dashed lines denote the theoretically ideal values in each plot.

The standard TD learning model without meta-learning showed healthy performance, *i*.*e*., good performance while switching targets, when the learning rate was the lowest (*α* = 0.1) (Fig. 3 black lines). As the learning rate increased, both the percentage of correct answers and the number of switches decreased rapidly. This was due to the fact that the decrease in *Q*-values associated with incorrect answers went back in time, *i*.*e*., the *Q*-values in the second and third states, where fixation on the central point should be maintained, also deteriorated, making it impossible to act according to the task event.

In the inverse temperature regulation (Fig. 3 blue lines), *β* was lowered from 100, which led to a greedy action choice, to 0.01, and yielded a more random choice when entering the exploration phase. Before the deterioration of *Q*-values in the second and third states where the fixation should be kept, the probability of losing concentration on the previous target increased, so the model exhibited healthy performance up to a higher learning rate than the standard model, but it still showed a decline in performance as the learning rate increased.

In contrast, the resetting of the *Q*-values avoided the deterioration of the *Q*-values in the second and third states even over a high learning rate because the gaze to the previous target was completely eliminated after one mistake. As a result, the results were close to the theoretically ideal values (success rate, 60% = 15/(1 + 9 + 15); number of target switch, 400 times = 10,000/(1 + 9 + 15): Fig. 3 dashed lines), assuming that the model made no mistake after it had found the correct answer through random search among the nine possible actions. These results indicate that resetting the value function is more effective than regulating the inverse temperature in meta-learning to achieve a quick transition from the exploitation phase to the exploration phase.

### 3.2 Performance of the NNRL model

We implemented a NNRL model with experience resetting (Fig. 2) and evaluated its performance. The model succeeded in learning the target search task after learning the center-fixation and fixed-target tasks without explicit mechanisms for search period judgement and meta-learning, as in the above TD learning models. In the first trial of the “first” fixed target task, the model was able to keep fixating on the central spot even when the peripheral spots were presented, while switching its gaze to a random direction after the central spot was turned off. After these random search trials, the model successfully gazed at the correct target. Conversely, on the first trial after a target was switched during the target search task, the model turned its gaze to the previous target. However, this behavior was reset after only one trial, and the model started searching randomly and reached the correct target within several trials. Also, note that the fixation on the central spot is not impaired in the second and third states. The success rate of the target–search task (Fig. 5A right) was sufficiently high, close to the ideal theoretical value of 60% mentioned above (Fig. 5A right dashed line).

**Fig. 4.**
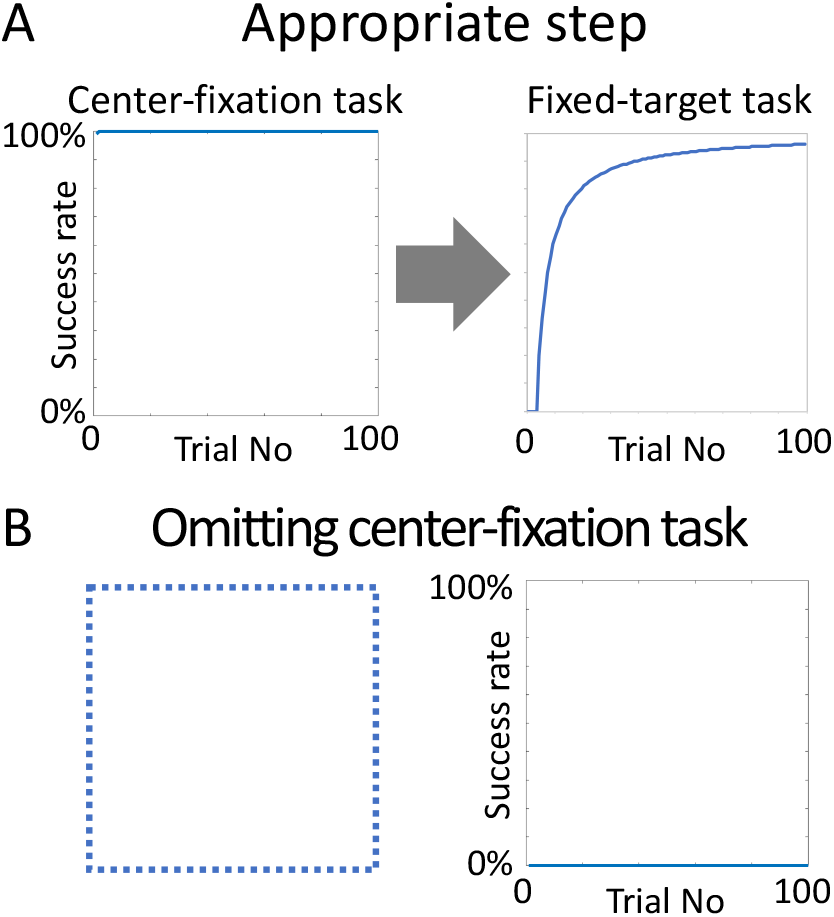
When learning of the fixation task was omitted, the fixed-target task could not be learned. (A) The success rate for each task when the appropriate step was taken. (B) The success rate of the fixed-target task when the fixation task was omitted. Success rates were calculated by moving average.

**Fig. 5.**
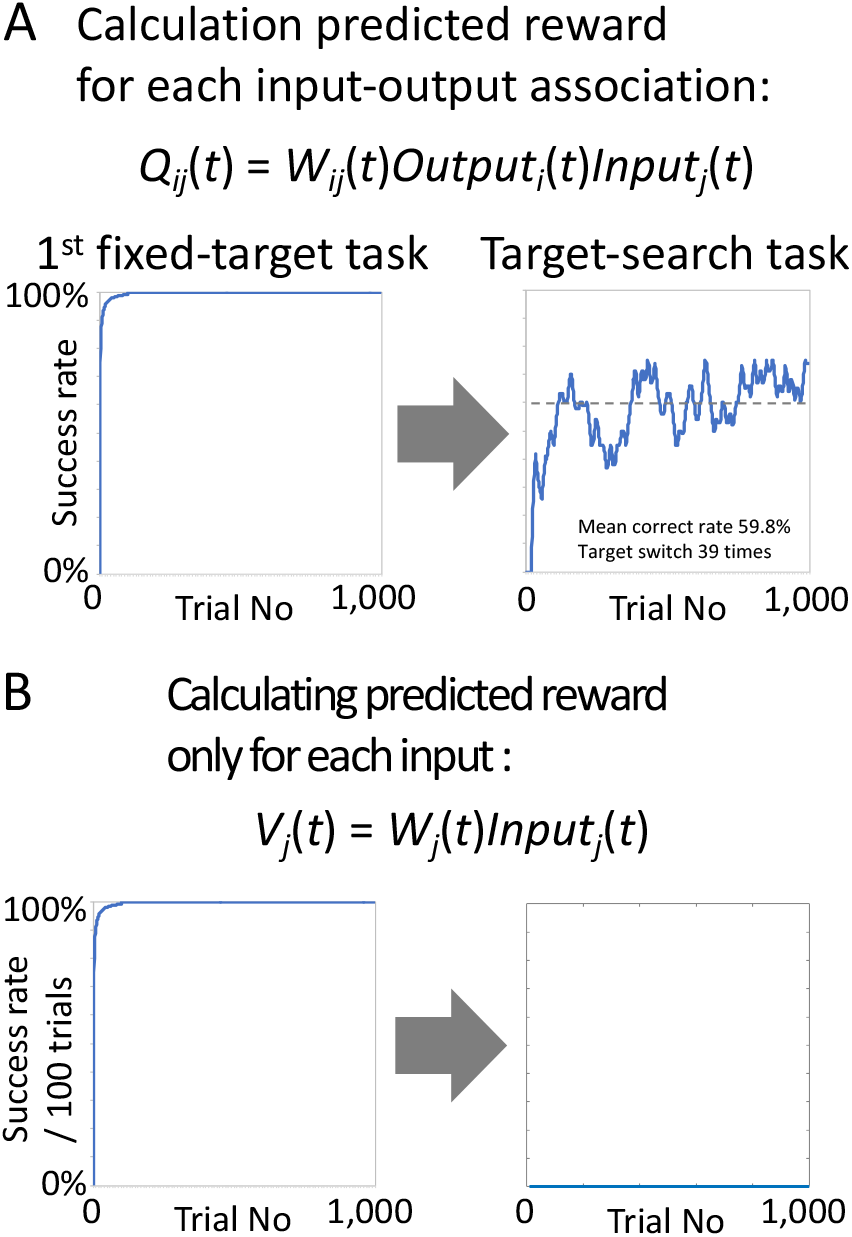
Comparison of learning ability among various modes of expected reward calculation. The success rate for each task when the expected reward was calculated for each input–output association (A) and only for each input (B). Success rates were calculated by moving average. In the target search task, the target was switched after 15 consecutive correct answers.

Learning needs to proceed via the appropriate steps. That is, if the center-fixation task is skipped, the model cannot learn the next fixed-target task (Fig. 4). At the stage of the fixed-target task, it was also important that the training for each target was completed with sufficient number of trials and that the training for all targets was also equal so that the contributions of the four spots to the output layer were equal. When the target search task was performed after the center-fixation task skipping fixed-target tasks, the number of trials for a particular target increased probabilistically. As a result, the appropriate experience resetting, *i*.*e*., the resetting of the input–output connection weights, did not occur when the target was switched.

For the NNRL model to modify the limited connection weights between the input and output layers in learning the target search task, it seems essential that reward prediction also be computed for specific input–output relationships. To test this idea, instead of *Q* as defined in the equation (6), the reward prediction *V* is calculated only for input *D* as follows:

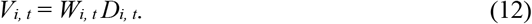

The target–search task was not learned at all (Fig. 5B right), even though the fixed-target task for the first target was learned correctly (Fig. 5B left). The training session using the fixed-target tasks was done for the four targets in sequence. The model using *V* failed to learn the second target, so it never succeeded in the target search task.

One of the advantages of combining neural networks with reinforcement learning is that it can be adapted to a variety of situations. In the target search task used for simulaitons in this study, there are 2^8^ = 256 variations in the presentation of peripheral spots. In the TD learning model above, it was necessary to design the *V*-table assuming that the variation in the presentation patterns will be available in advance. By contrast, the NNRL model did not require such a preliminary design, and was able to learn a target search task for various spot patterns by following the appropriate training procedure, shown in Fig. 1B. An example is shown in Fig. 6, where the correct response rate and the number of switches were comparable to the theoretically ideal values (Fig. 6 dashed lines), even when the number of spots was increased by placing them in arbitrary positions.

**Fig. 6.**
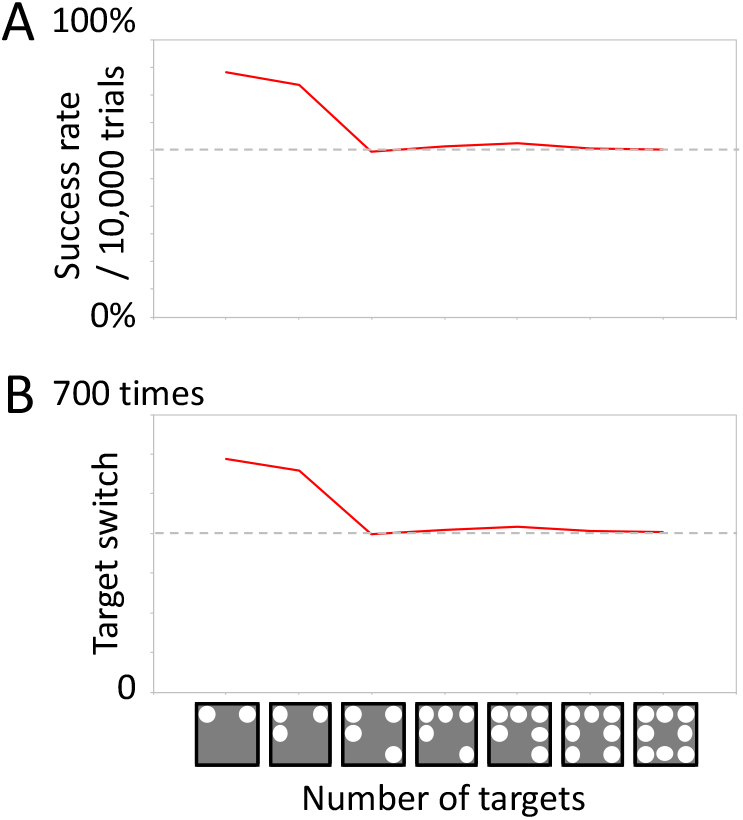
Demonstration of the flexible performance of the NNRL model for different numbers of arbitrarily placed spots. The learning rate was set to the default value (*α* = 0.25). The success rates (A) and the number of target switches (B) in the target search task after the appropriate pre-lessons. Each dashed line is a theoretically ideal value of the random search among nine possible actions during the exploration phase.

Interestingly, when the number of targets was two or three, the results were better than the theoretically ideal values of random search among nine possible actions. This was due to the fact that the learning rate was set to 0.25, which means that the reset was not completely applied. That is, because of rectification of *E* defined in equation (9), the relevant input–output connection weights converged to 1/2 or 1/3 (> 0.25) in these cases, which caused a random search only among the target candidates. In fact, the model showed almost the same values as the theoretically ideal values in the case of random search among the target candidates (for two targets, the correct response rate was 15/(1 + 1 + 15) = 88%, and the number of switches 10,000/(1 + 1 + 15) = 588 times; for three targets, the correct response rate was 15/(1 + 2 + 15) = 83%, and the number of switches 10,000/(1 + 2 + 15) = 556 times). However, if we lowered the learning rate sufficiently (*e*.*g*., to 0.1 < 1/8) so that the random search among target candidates would happen for all possible cases, a trade-off emerged: learning would not proceed as well in such a case.

## 4 Discussion

The exploration–exploitation trade-off problem is a major challenge in reinforcement learning for both animals and machines. In performing a task in which the exploration phase and the exploitation phase are switched back and forth, accurately recognizing the exploration phase and increasing exploratory behaviors by meta-learning will lead to improved performance in reinforcement learning models. To increase exploratory behavior, we showed that resetting the experience is a better meta-learning method than is reducing the greediness of action selection, which was proposed in a previous study [4]. Furthermore, as a step toward future construction of a model more closely resembling the brain, we developed a prototype NNRL model that resets the experience in a biologically plausible way compared to TD learning models, and illustrated that it can adapt to various situations.

In this study, to represent various input–output relationships with a small number of units, we used a two-layer neural network (Fig. 2A) and developed a model with a dual structure in which the relationship between the input and output layers was modified by the reward prediction error calculated by a separate subsystem (Fig. 2B). This model’s dual structure is orthodox for the integration of neural networks and reinforcement learning, and it is expected that greater flexibility can be achieved by adding more layers to the two subsystems in the future [5, 6]. In addition, the fact that a reward prediction error with a small number of degrees of freedom (9 in this model) modulates a network with a large number of degrees of freedom (9^2^ in this model) is consistent with the fact that in the real brain, a small number of dopamine neurons, which are thought to encode the reward prediction error, project widely in the cerebrum and are involved in modulating the network [7]. In the future, more realistic models will help us to understand the real brain activity. In this sense, the model is promising, although it still needs improvement in terms of flexibility to changes in the temporal structure of behavioral tasks.

Whether the value function should be calculated for the combination of input and output or only for the input, *i*.*e*., which action value function *Q* or state value function *V* should be used, remains controversial. As shown in Fig. 5, our NNRL model clearly needed to modulate only a limited number of input-to-output connection weights to learn our target search task. However, when we computed the value function for a combination of input and output, our TD learning model fell into a stable state where the model alternated between wrong and right answers without switching targets (data not shown), whereas, when we obtained the value function from the input only, it showed preferable performance (Fig. 3). In fact, the basal ganglia, which plays an important role in reinforcement learning in the brain, is thought to contain not only a state value function but also an action value function [8]. Furthermore, a model has been proposed in which the two functions are used flexibly depending on how much control the agent’s actions have over the outcome [9]. The real picture is that two value functions exist in parallel, and the dominant value function is determined by which one yields more unique predictions.

In the first place, primates not only reset predictions that are no longer valid but also switch to other predictions. Monkeys do not relearn from scratch every time the task situation changes [10,11]. Some parts of the mechanism of learning and behavioral switching may have appeared in the cases of two- and three-target candidates in Fig. 6, but this is an important issue to consider to develop this model into one that is more general and brings about a better understanding of the brain’s structure and functions.

## Acknowledgements

This work was supported by JSPS KAKENHI Grant Number JP16H06276 (Platform of Advanced Animal Model Support), 17K07060, 20K07726 (Kiban C), MEXT KAKENHI Grant Number 20H05478 (Hyper–Adaptability) and Japan Agency for Medical Research and Development (AMED) under Grant Number JP18dm0207051.

The authors declare no competing financial interests.

